# The DNA end-binding protein Ku associates with human telomeres primarily via protein-protein interactions

**DOI:** 10.1101/2019.12.11.873422

**Authors:** Ann T. Sukumar, Christopher L. Williams, Celina Y. Jones, Elif Asik, Danna K. Morris, Alessandro Baldan, Sandra M. Indiviglio, Ilaria Chiodi, Chiara Mondello, Alison A. Bertuch

## Abstract

The Ku heterodimer (Ku70/Ku80) binds DNA ends with high affinity but without sequence specificity and, upon binding ends created by double-stranded breaks (DSBs), initiates canonical nonhomologous end-joining (c-NHEJ). Ku also localizes to functional telomeres where its c-NHEJ activity is inhibited. Interestingly, Ku has been co-opted at telomeres across species, where it performs varied telomeric functions. In humans, Ku is essential for its role in telomere maintenance, but how it associates with human telomeres is not known. Analysis of Ku’s telomere association in different populations of cen3tel cells, which had a wide range of average telomere lengths, supported Ku’s localization at human telomeres primarily via protein-protein interaction. We also found that the Ku70 and Ku80 α5 helices, which are on opposing sides of the heterodimer and were previously implicated in *Saccharomyces cerevisiae* Ku’s NHEJ and telomeric functions, respectively, participated in Ku’s telomere association in human cells. While the Ku70 α5 mutant showed increased interaction with TRF2, the Ku80 α5 mutant was not impacted for TRF2 association. Interestingly, residues altered to impair Ku’s DNA end-binding function were also involved in TRF2 interaction and telomere association. Overall, our results suggest protein-protein interactions as the primary mode by which Ku associates with human telomeres.

## INTRODUCTION

Genomic integrity is preserved when DNA double-strand breaks (DSBs) produced by both endogenous and exogenous factors are dutifully repaired. Two main pathways govern the repair of DSBs, namely classical (aka canonical) non-homologous end joining (c-NHEJ) and homologous recombination (HR). Each of these pathways employs a host of protein factors to identify and process broken ends in a coordinated fashion, which ultimately culminates in repair (1).

To maintain genomic stability, the natural ends of eukaryotic chromosomes, however, must be concealed from DNA repair machineries, as they have the potential to appear as DSBs and be used as substrates for c-NHEJ or HR. To prevent this mishap, chromosome ends take the form of specialized DNA-protein structures, known as telomeres, that inhibit recognition of chromosome ends as DSBs (2). Mammalian telomeric DNA is comprised of tandem repeats of TTAGGG sequence, which spans several kilobases in length, terminating in a 3’ overhang. A complex of six proteins, collectively known as shelterin, is found along the telomeric tract (3). Shelterin contributes to telomere length homeostasis by facilitating the recruitment and function of telomerase, a reverse transcriptase that replenishes telomeric repeats lost each cell division cycle due to the inability of the semi-conservative replication machinery to faithfully replicate the chromosome end (4). Whereas cells can divide for numerous generations in the absence of telomerase, the presence of shelterin is essential, as it blocks improper access of proteins involved in DNA repair at telomeres and, thus, functions to maintain genomic stability (5).

Specifically, TRF2, a component of the shelterin complex that binds the duplex telomeric repeats, is crucial to counteract c-NHEJ at telomeres. Telomeres lacking TRF2 display extensive c-NHEJ-mediated chromosome end-to-end fusions (6-8). One mechanism by which TRF2 is thought to impede c-NHEJ is by enabling the formation of the t-loop structure at the telomeric DNA end, wherein the terminal single stranded 3’ overhang engages the upstream double stranded telomeric DNA via strand invasion, thereby masking the chromosome ends within the t-loop (9,10).

Despite the masking of telomere ends within the t-loop structure, the c-NHEJ factor Ku associates with functional telomeres (11). This evolutionarily conserved protein, comprised of the Ku70 and Ku80 subunits, is well-studied for its role in promoting c-NHEJ. In addition to stabilizing each other upon heterodimerization, the two subunits form a ring-shaped channel through which Ku can efficiently associate with DNA ends irrespective of sequence (12,13). Due to its abundance and high affinity for DNA ends, Ku binds DSBs within seconds and sets in motion the molecular events necessary to achieve end joining (14).

While the configuration of the t-loop might prohibit Ku from readily loading onto the telomeric DNA termini, disassembly of the t-loop presumably occurs to permit telomere replication and this could then provide access for Ku to the telomeric end. Studies on *Saccharomyces cerevisiae* Ku mutants impaired for DNA end-binding, but not for heterodimerization nor interaction with other known telomere-related factors, revealed that Ku’s end-binding function is pivotal for telomere length maintenance, telomere end protection, and telomeric silencing in that species (15). However, it is yet to be determined if Ku associates with human telomeres by loading onto the DNA ends, and, if so, whether direct loading is required for telomeric integrity in human cells. In addition to TRF2, Ku has been shown to interact with the shelterin proteins TRF1 and Rap1, but the functional consequences of these associations and whether Ku associates with mammalian telomeres solely via protein-protein interactions are not clearly understood (16-18).

Furthermore, how Ku is restricted from engaging in c-NHEJ at telomeres is also not well understood. Structural studies highlight Ku’s polarity when situated at the DNA end with the α/β domain of Ku70 facing toward the DNA end and α/β domain of Ku80 on the opposite side, distal from the end (12). Follow-up studies based on analyses of separation-of-function yeast Ku mutants suggested that, when Ku loads onto a DNA end, residues pertinent to Ku’s c-NHEJ functions are located outward towards the DNA end whereas residues associated with Ku’s telomeric functions are positioned inward distal from the DNA end (19). This functional organization of Ku might allow Ku’s c-NHEJ functions to be suppressed at telomeres while keeping its telomeric functions intact. Evidence for this line of thought came from the examination of a Ku70 α5 mutant, which harbors mutations in the conserved α-helix 5 and has been found to be defective in DNA repair in both yeast and mammalian cells (19-21). Particularly, in human cells, evidence suggests that Ku70 α5 may participate in TRF2 interaction as well as in bridging DNA ends at DSBs via Ku-Ku heterotetramerization. This finding implies that Ku’s association with TRF2 could obstruct its propensity to bridge DNA ends and, thereby, inhibit c-NHEJ at telomeres (21).

Interestingly, Ku is involved in various aspects of telomere length maintenance across many species. For instance, loss of either the Ku70 or Ku80 subunit in *S. cerevisiae* results in stably short telomeres (22,23). While inconsistencies exist with respect to how Ku influences telomere length in murine cells, with studies showing both positive and negative regulation of telomere length in Ku knockout mice (24,25), reduction or loss of Ku in human cells results in substantial telomere loss (26,27). In cells with conditional knockout of Ku80, this is associated with an increase in telomeric t-circles, which have been proposed to have formed due to HR de-repression at telomeres (26). Cells deficient for Ku in other species, including yeast and mice, also exhibit aberrant HR-mediated telomere phenotypes, suggesting that, in addition to inhibiting HR at DSBs, an important function of Ku at telomeres is to prevent deleterious recombination events (7,28-31). Importantly, Ku is not an essential gene in budding yeast or mice, but it is required for viability in human cells, particularly for its critical role in preventing severe telomere loss (26,32-34)

Contrary to *S. cerevisiae*, where probing separation-of-function Ku mutants have yielded valuable knowledge regarding Ku’s association and multifaceted functions at telomeres, not much is known about how Ku functions at human telomeres. Given Ku’s ability to readily bind DNA ends to initiate c-NHEJ and its significance in maintaining functional telomeres to confer viability, an outstanding question in the field is how Ku associates with human telomeres. In this study, we interrogated Ku’s association with human telomeres using a unique cell line, cen3tel, that developed hyper-elongated telomeres during propagation in culture(35) and also using a system in which endogenous Ku was depleted while mutant Ku was expressed at endogenous levels. Alpha helices on opposing sides of the heterodimer and cognate to helices that mediate NHEJ or telomeric functions in budding yeast were mutated, as were residues involved in Ku-DNA interaction. Analysis of Ku’s association with telomeres in cen3tel cells at different stages of propagation, and thus characterized by telomeres of different length, was consistent with Ku associating with telomeres primarily via protein-protein interactions. Consistent with this, we found that mutations in the Ku70 and Ku80 α5 helices impacted Ku’s recruitment to human telomeres as did mutations that impacted DNA end-binding. These human Ku mutants were differentially affected for association with TRF2, with the Ku DNA end-binding mutant showing a reduced association and the Ku70 α5 mutant, which is known to impact NHEJ, showing increased association. Our results lead us to propose that Ku associates with human telomeres primarily via interaction with telomere-bound proteins.

## MATERIALS AND METHODS

### Cell culture and siRNA transfections

Cen3tel cells were cultured in DMEM supplemented with 1X L-Glutamate, 1X non-essential amino acids, and 10% FBS. Flp-In T-REx 293 cells (Invitrogen) were cultured in DMEM media containing 10% FBS, blasticidin (15 μg/mL, Invitrogen), and Zeocin (100 μg/mL, Invitrogen). To integrate the transgenes, 6X N-terminally Myc-tagged cDNAs encoding Ku70 or Ku80 WT and mutants were cloned into pcDNA/FRT/TO (Invitrogen) and then transfected along with Flp recombinase expressing pOG44 plasmid using Lipofectamine transfection reagent (Invitrogen) as per the manufacturer’s instructions. Stably transfected positive integrants were selected in media containing blasticidin and hygromycin B (100 μg/mL, Invitrogen).

For siRNA transfections, cells were plated in 6 well plates and transfected with 5 μl Lipofectamine RNAiMAX transfection reagent and siRNAs (Invitrogen) at 50 nM final concentration according to the manufacturer’s instructions. To deplete endogenous Ku70, pooled siRNAs targeting 3’ UTR were used (5’-CCCACUUUGCUGUUCCUUATT-3’, 5’-CCUGAAUAAAGAGCCCUAATT-3’, 5’-CAUAAGUCGAGGGACUUUATT-3’). The siRNA used to knockdown endogenous Ku80 was as follows: 5’-GCGAGUAACCAGCUCAUAATT-3’. Silent mutations were introduced in the Myc-tagged Ku80 WT and Ku80 mutant cDNAs to prevent siRNA recognition. At 24 hours following siRNA transfection, cells were trypsinized and plated with media containing 1 μg/mL doxycycline (Sigma-Aldrich).

### Chromatin immunoprecipitation

Chromatin immunoprecipitation assays were performed as described previously (40), except to immunoprecipitate endogenous Ku80 in cen3tel cell lines, 7 μl of anti-Ku80 antibody (2753, Cell signaling) was used. To immunoprecipitate Myc-Ku70, 3.5 μl anti-c-Myc antibody (M4439, Sigma-Aldrich) and 50 μl Pierce protein G magnetic beads (88848, Thermo Fisher Scientific) were used and to immunoprecipitate Myc-Ku80, 50 μl of Pierce anti-c-Myc magnetic beads (88843, Thermo Fisher Scientific) was used for each sample.

### Electrophoretic mobility shift assay (EMSA)

For EMSA, cells were lysed in lysis buffer (50 mM Tris-HCl pH 7.5, 1 mM EDTA, 400 mM NaCl, 1% Triton X-100, 0.1% SDS, 1 mM dithiothreitol [DTT], 1 mM phenylmethylsulfonyl fluoride [PMSF], and 1X protease inhibitor cocktail III [Calbiochem]). For the binding reaction (20 μl), 5 μg of each sample was incubated with 1.5 ng ^32^P end-labeled DNA fragment (198 bp) in binding buffer (20 mM Tris-HCl pH 8, 150 mM NaCl, 10 mM EDTA and 10% glycerol) for 30 minutes at room temperature, together with 1 μg closed circular pcDNA3.1. Following addition of 0.5 μl of 5X bromophenol blue, the samples were loaded onto a 6% polyacrylamide gel and electrophoresed in Tris-glycine buffer at 45 mA for approximately 1 hour. The gel was then exposed to a phosphorimager screen and imaged using Storm865 imaging system (Molecular Dynamics).

### Co-immunoprecipitation

Cells for each sample were harvested from a 10 cm plate. Following two washes in 1X phosphate-buffered saline (PBS), cells were subjected to lysis in 250 μl lysis buffer (50 mM Tris-HCl at pH 7.5, 1 mM EDTA, 150 mM NaCl, 1% Triton X-100, 1 mM PMSF and 1X protease inhibitor cocktail III [Calbiochem]). After a 5-minute incubation on ice, 12.5 μl NaCl was added, and following another 5-minute incubation, 250 μl ice cold water was added. The lysates were then clarified at 4 °C and 14,000 rpm for 10 minutes, and the protein concentration was quantified using the BCA protein assay kit (Pierce). For Myc immunoprecipitations, either 3.5 μl anti-c-Myc antibody (M4439, Sigma-Aldrich) and 50 μl of Pierce protein G magnetic beads (88848, Thermo Fisher Scientific) or 50 μl of Pierce anti-c-Myc magnetic beads (88843, Thermo Fisher Scientific) was used. To immunoprecipitate TRF2 and for IgG control, 4 μl of anti-TRF2 (NB110-57130, Novus Biologicals) and 4 μl of rabbit IgG (2729, Cell Signaling) were used, respectively. DNase treatment of indicated samples was carried out by incubation of each sample with 10 μl of DNaseI (NEB), 5 μl of Benzonase Nuclease (Novagen) and MgCl_2_ (1 mM final concentration) for 1 hour on ice. For each experiment, 500-600 μg of lysate was immunoprecipitated overnight at 4 °C on a rotator and the beads were washed four times in lysis buffer. The immunoprecipitates and 5% inputs were resolved on 10% SDS-PAGE gels and transferred to Immobilon-FL PVDF membranes (Millipore). The following antibodies were used to probe the membranes: anti-TRF2 (NB110-57130, Novus biologicals, 1:2000 dilution), anti-Ku80 (sc-9034, Santa Cruz, 1:1000 dilution; sc-5280, Santa Cruz, 1:250 dilution), anti-Ku70 (MS-329-P1, Lab vision, 1:1000 dilution), anti-Rap1 (A300-306A-2, Bethyl, 1:1000 dilution), anti-c-myc (M4439, Sigma-Aldrich, 1:2000 dilution), and anti-β-actin (A5441, Sigma-Aldrich, 1:5000 dilution). To strip and re-probe, the membranes were washed in 0.4 M NaOH for 15 minutes.

### Telomere length analysis

One million cells per sample were incorporated in 1% agarose plugs (Bio-Rad) and the genomic DNA was subjected to digestion with HinfI (NEB) and RsaI (NEB) and RNase treatment overnight. The agarose plugs were then loaded onto a 1% agarose gel and electrophoresed in 0.5% Tris-borate-EDTA in a pulse-field apparatus (Bio-Rad) at 14 °C (6 V/cm, 13 hours, 5 seconds initial and 20 seconds final). The gel was washed in depurination buffer (0.25 M HCl) for 20 minutes, denaturation buffer (0.5 M NaOH, 1.5 M NaCl) for 30 minutes, and neutralization buffer (0.5 M Trish pH 7.5, 3 M NaCl) for 30 minutes and the DNA was transferred to a Zetaprobe membrane (Bio-Rad) in 20X SSC. The membrane was UV crosslinked, blocked in hybridization buffer (0.5 M phosphate buffer, pH 7.2, 1 mM EDTA, 7% SDS) for 2 hours at 65 °C, and hybridized overnight to an 800 bp telomere probe randomly labeled with ^32^P-dCTP. Following washes in low stringency (4X SSC, 0.1% SDS) and high stringency buffers (2X SSC, 0.1% SDS), the membrane was subjected to phosphorimager scanning. Telomere length was quantified using TeloTool. The mean telomere length is designated as a red dot (41).

### 2D gel electrophoresis

Cells were harvested 4 days after siRNA transfection, and DNA was extracted using Qiagen’s DNeasy kit. Samples were digested overnight at 37 °C in HinfI, RsaI, and RNase, then ethanol precipitated. Eight to 10 μg of digested DNA was loaded in a 10 x 12 cm 0.4% agarose gel lacking ethidium bromide (EtBr) and was run in the first dimension at 1 V/cm for 15 hours at room temperature. The gel was stained in 0.3 μg/mL EtBr for 30 minutes at room temperature; then the lanes were excised and placed 90° counterclockwise in a 20 x 25 cm tray. A 1% gel with 0.3 μg/mL EtBr was cast around the lanes. Electrophoresis in the second dimension was performed at 5 V/cm for 8 hours at 4°C. The gel was washed in depurination buffer (0.25 M HCl) for 20 minutes, denaturation buffer (0.5 M NaOH, 1.5 M NaCl) for 30 minutes, and neutralization buffer (0.5 M Trish pH 7.5, 3 M NaCl) for 30 minutes. It was transferred to a nylon membrane (Bio-Rad Zeta-probe), UV crosslinked, and the membrane was pre-hybridized in hybridization buffer (0.5 M phosphate buffer pH 7.2, 1 mM EDTA, 7% SDS) for at least 2 hours at 65 °C. It hybridized overnight at 65 °C to an 800 bp telomere probe randomly labeled with ^32^P-dCTP. Excess probe was washed off, and the signal was detected by phosphorimager scanning.

### Annexin V staining

Annexin V staining assays were performed as per the manufacturer’s instructions (BioLegend). Briefly, cells were harvested 4 days after siRNA transfection, washed twice with PBS, and resuspended in Annexin V binding buffer (BioLegend). Cells were then stained with FITC Annexin V and PI for 30 minutes and the percentage of apoptotic and viable cells was analyzed by flow cytometry (BD LSRII).

## RESULTS

### Ku associates with human telomeres principally by protein-protein interactions

To investigate whether Ku associates with telomeres principally by loading onto the telomere end via its DNA end-binding activity or by protein-protein interaction, we first capitalized on the cen3tel cell line, which was derived from a centenarian’s skin fibroblasts following immortalization by hTERT retroviral infection (35,36). Notably, after an initial period of telomere shortening, the cen3tel line progressively developed hyper-elongated telomeres over successive population doublings (PDs) resulting in telomere lengths ranging from 10 kb (around PD 150) to as long as 100 kb (around PD 1000) (35). Thus, cen3tel cells collected at various PDs allowed us to analyze Ku’s telomere association in cells with a wide range of telomere lengths while derived from the same source. We reasoned that, if Ku associates with telomeres principally via DNA end-binding, the amount of telomeric DNA immunoprecipitated with Ku80 in chromatin immunoprecipitation (ChIP) assays would be equivalent across different PDs of the cen3tel cell line with a wide range of telomere lengths because the number of telomeric ends would be comparable across PDs (Fig. 1A). Because the amount of Alu DNA, used as a repetitive DNA control, would also be similar across the different PDs, the ratio of telomeric DNA immunoprecipitated with Ku80 to Alu DNA in the input would remain unchanged across PDs with increasing telomere lengths. In contrast, because the amount of telomeric DNA in the input would be higher in successive PDs as telomere length increased, the ratio of immunoprecipitated telomeric DNA to input telomeric DNA would decrease from low PD to high PD with increasing telomere length. On the other hand, if Ku associates with telomeres principally through protein-protein interactions along the telomere tract, the ratio of immunoprecipitated telomeric DNA to input Alu DNA would increase with increasing telomere lengths, whereas the amount of telomeric DNA immunoprecipitated with Ku80 as the telomere length increased would either remain the same, if the distribution of binding partners remained unchanged, or decrease, if the distribution of the binding partners decreased across the telomeric tract, e.g., as observed for chromatin-bound TRF2 and RAP1 in HeLa subclones with long as compared to short telomere lengths (37).

**Figure 1.**
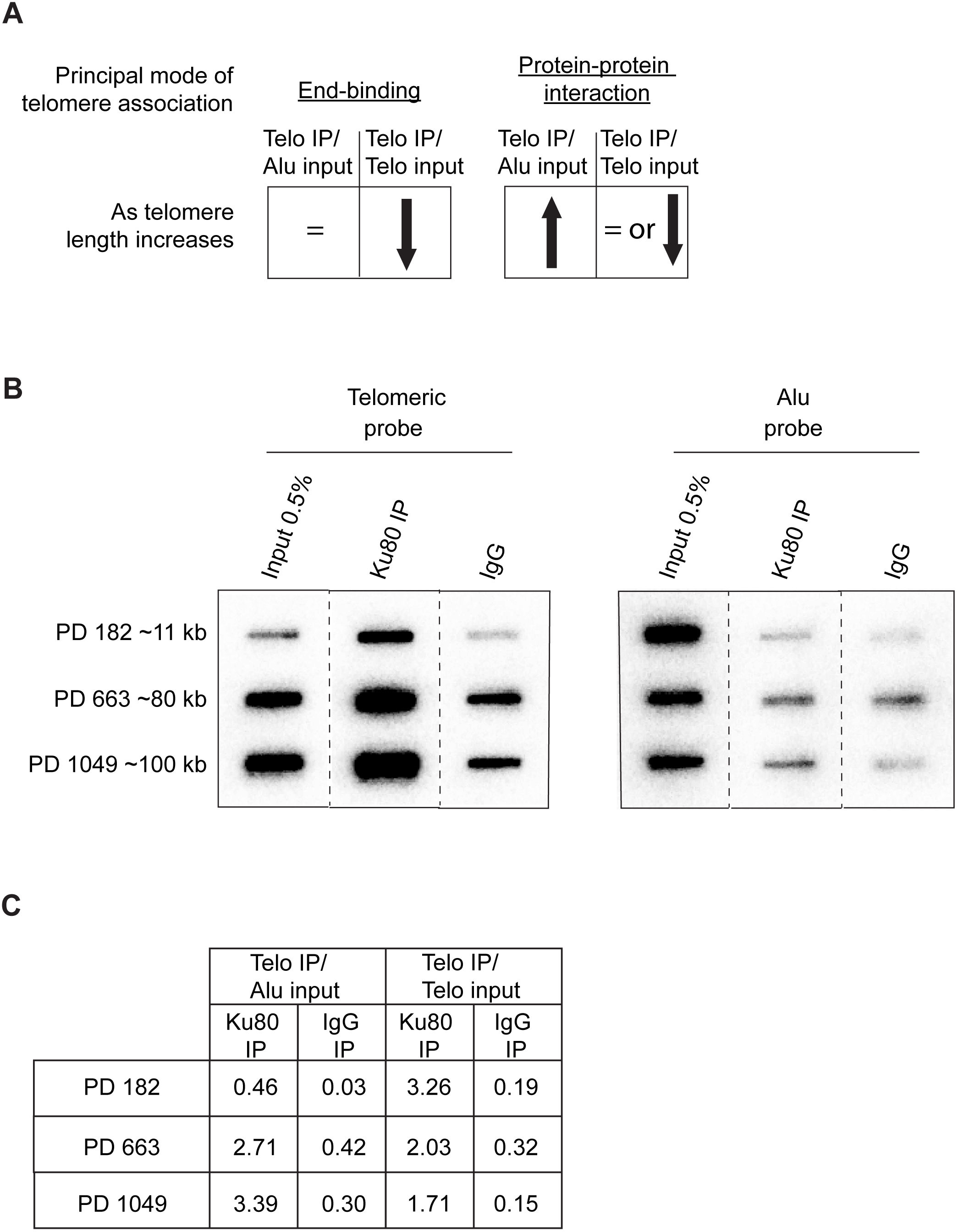
Ku localizes to telomeres via protein-protein interactions. **(A)** Expected trends as telomere length increases across cen3tel PDs for the quantity of telomeric DNA in the Ku80 chromatin immunoprecipitation ChIP (Telo IP) relative to either the quantity of Alu DNA or telomeric DNA in the input (Alu input or Telo input, respectively) based on whether Ku’s principal mode of association with telomeres is either via end-binding or protein-protein interactions. **(B)** Representative ChIP of cen3tel cells pertaining to the designated PDs with the approximate telomere lengths. Cells were crosslinked, sonicated, and subjected to lysis followed by immunoprecipitation with a Ku80 antibody. Telomeric and Alu DNA associated with the respective PDs was analyzed via slot blot. IgG was used as negative control and Alu DNA was used as repetitive DNA control. **(C)** Table showing the percentage of telomeric DNA immunoprecipitated with Ku80 or control IgG relative to telomere input and Alu input for the respective PDs. Additional representative experiments at slightly different PDs are shown in Supplementary Figure S1.

We performed Ku80 ChIP assays using cen3tel cells that were collected at low, mid and high PDs, harboring ∼11 kb, 80 kb, and 100 kb telomere lengths, respectively (Fig. 1B). Due to the repetitive nature of telomeric repeats, the presence of telomeric DNA in the immunoprecipitates was assayed by dot blotting. Quantification of ChIP data revealed clear evidence for Ku associating with telomeres via protein-protein interactions as we detected an increase in the immunoprecipitated telomeric DNA to input Alu DNA ratio with increasing telomere length (Fig. 1C and Supplemental Fig. S1). We also observed a decrease in the ratio of immunoprecipitated telomeric DNA relative to input telomeric DNA from low to high PD, which could be consistent with either mode of interaction. However, as the length of telomeric DNA increased almost 10-fold from low PD to high PD (from 11 kb to 100 kb) and the ratio of telomeric IP/telomeric input only decreased approximately 2-fold, the results are most suggestive of Ku primarily associating with telomeres indirectly via protein-protein interactions. Nonetheless, we cannot exclude the possibility of direct engagement of the ends at some or even all telomeres.

### Experimental system and initial characterization of Ku mutants used to probe the role of Ku70 α5, Ku80 α5, and Ku’s DNA end-binding activity at telomeres

Given the apparent role of protein-protein interactions for Ku’s association with human telomeres, we next sought to elucidate if the structural and functional organization of Ku that we identified in yeast (19) is conserved in humans by investigating the impact of mutations in the Ku70 and Ku80 α5 helices on Ku’s telomere association and function (Fig. 2A). These helices localize to the Ku70 and Ku80 α**/**β domains, which are thought to provide sites for interactions with protein binding partners (12). We also investigated the impact of mutations designed to impair DNA end-binding, which would allow us to further probe whether Ku associates with functional telomeres via direct end-binding. We engineered the Flp-In T-REx 293 cell line to express Myc-tagged wild type (WT) or mutant Ku70 or Ku80 transgenes under the control of the doxycycline-inducible promoter (Fig. 2B). Our experimental approach entailed depletion of endogenous Ku70 or Ku80 using siRNA, followed by induction of the integrated constructs by doxycycline addition, and, 72 hours later, harvest of cells for functional analyses (Fig. 2C). Endogenous Ku70 was depleted via siRNAs targeting the 3’ UTR, and siRNA-resistant Myc-Ku80 transgenes were integrated to mask the recognition by Ku80 siRNA. Using this strategy, we achieved robust knockdown of endogenous Ku70 and Ku80 and, importantly, expression of the corresponding transgenes at levels comparable to endogenous, thereby, allowing us to test the mutants in a system in which exogenous Ku was not massively overexpressed (Figs. 2D-E).

**Figure 2.**
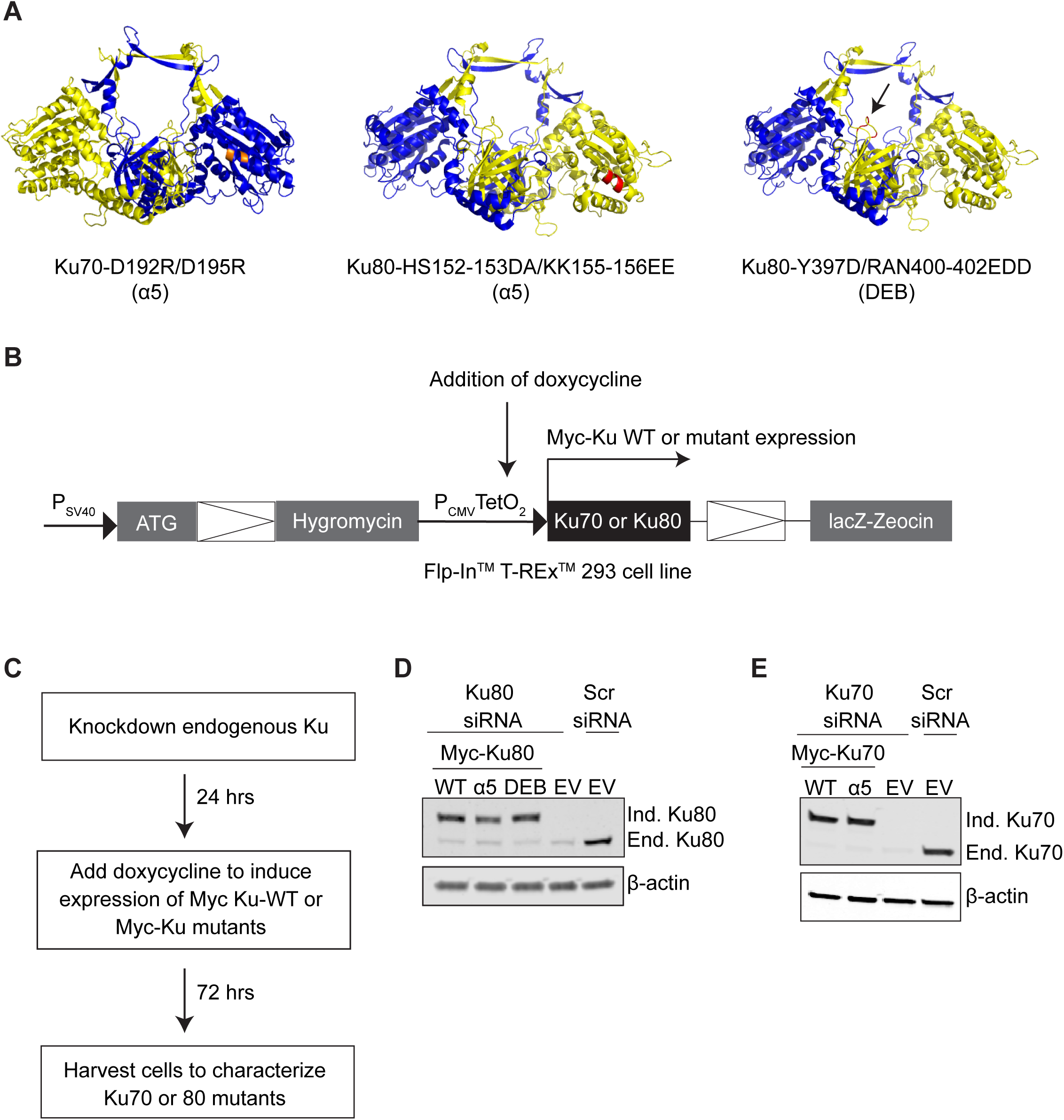
Schematic of human Ku mutants analyzed in this study and system for depleting endogenous Ku70 or 80 and expressing wildtype and mutant transgenes. **(A)** Cartoon representation of the crystal structure of human Ku bound to DNA (PDB 1JEY(12)) showing Ku70 in blue and Ku80 in yellow. Residues in Ku70 α5 and Ku80 α5 mutated in this study are in orange and red, respectively. Residues mutated to generate Ku80-DEB are shown in red and indicated by the arrow. DNA is hidden. Specific amino acid substitutions for each mutant is indicated below the corresponding structure. **(B)** Schematic of integrated expression construct in Flp-In T-Rex™ 293 cell line. Doxycycline-inducible promoter controls expression of Myc-tagged WT or -mutant Ku70 or Ku80 transgenes. **(C)** Experimental strategy. **(D)** Western blots showing Ku80 protein levels following siRNA mediated depletion of endogenous Ku80 and induction of the indicated Ku80 transgenes or empty vector. **(E)** Same as **D** except showing Ku70 protein levels and induction of the indicated Ku70 transgenes or EV. Scr, scrambled siRNA; WT, wild type; EV, cell line with an empty vector integrated; Ind, induced; End, endogenous; The transgenes migrate at higher molecular weights than endogenous Ku due to the N-terminal myc tags. β-actin was used as loading control.

The Ku70 α5 mutant transgene bore mutations in residues cognate to those required for c-NHEJ, but not telomeric functions, in yeast and the same mutations we previously demonstrated to impact DNA repair in mammalian cells (21) (Fig. 2A left). The residues mutated in the Ku80 α5 mutant transgene were cognate to those required for interaction with Sir4, which is important for telomeric silencing and telomere length maintenance, but not c-NHEJ, in yeast (19,38) (Fig. 2A middle). Lastly, the residues mutated in the Ku80 DEB transgene, which localized to the DNA binding channel, were similar to those that impaired DNA end-binding when mutated in yeast (15) (Fig. 2A right).

As an initial characterization, we performed electrophoretic mobility shift assays (EMSAs) to assess DNA end-binding using whole-cell extracts (WCEs) prepared from cells in which endogenous Ku70 or Ku80 was depleted and either Myc-WT or Myc-mutant transgene of the corresponding gene was expressed. The Ku70 α5 mutant shifted the radiolabeled fragment similarly to WT Ku70 indicating that the Ku70 α5 mutant was capable of DNA end-binding (Fig. 3A). Thus, the previously reported DNA repair defect resulting from the Ku70 α5 mutation was not a consequence of impaired DNA end-binding (21). Similar EMSAs results were obtained when endogenous Ku80 was depleted and either Myc-Ku80 WT or Myc-Ku80 α5 mutant was expressed, indicating any defects detected with the Ku80 α5 mutant could not be attributed to defective end-binding. In contrast, WCEs expressing Myc-Ku80 DEB failed to shift the radiolabeled DNA suggesting that the residues mutated in Ku80’s DNA loading channel indeed rendered Myc-Ku80 DEB defective for DNA end-binding.

**Figure 3.**
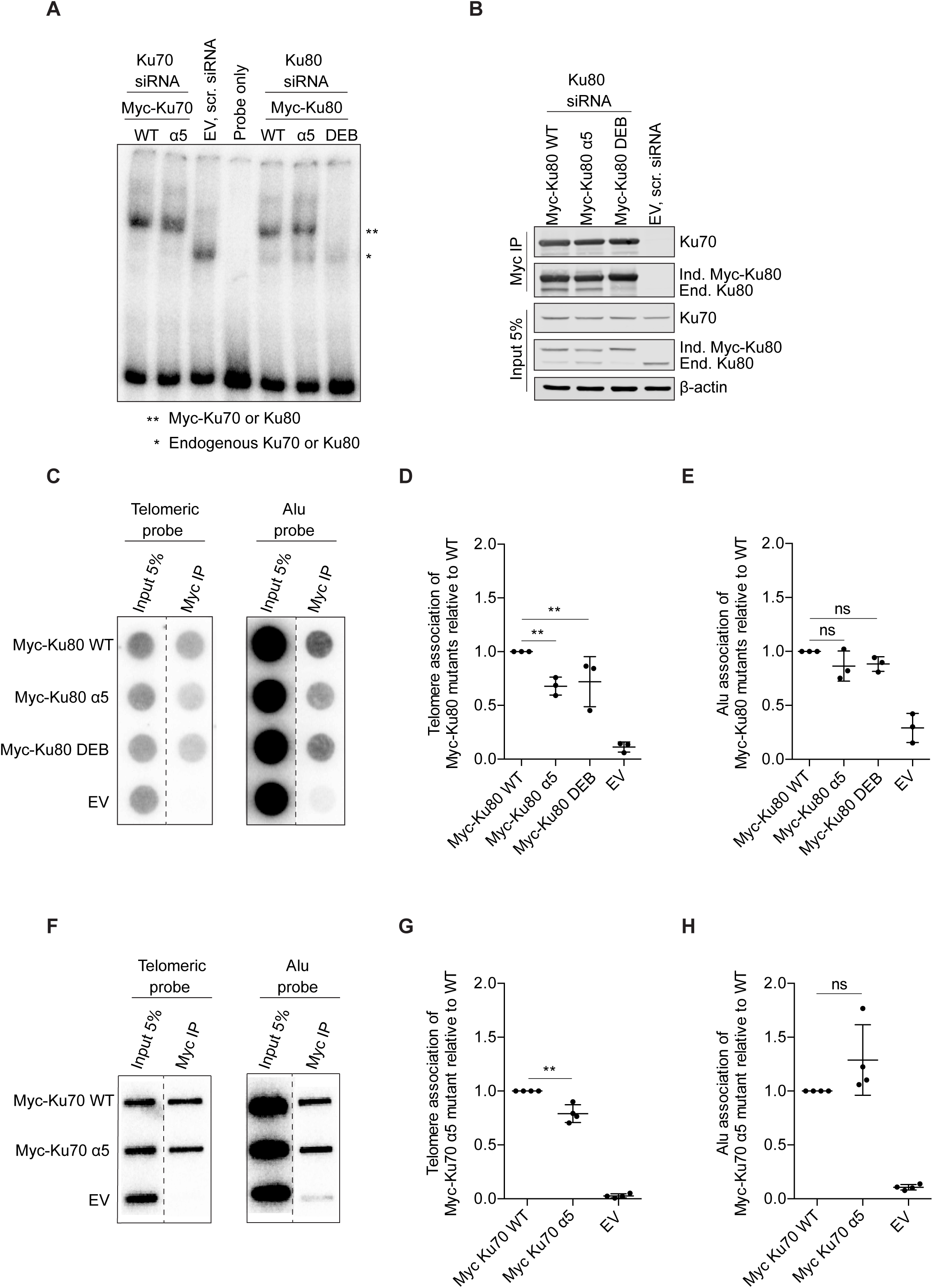
Myc-Ku80 α5 mutant and Myc-Ku80 DEB exhibit decreased association with telomeres. **(A)** Electrophoretic mobility shift assays (EMSA) using whole cell extracts (WCE) from cells induced to express Myc tagged WT or mutant Ku70 or Ku80 or EV following siRNA knockdown of endogenous Ku70 or Ku80. WCE were incubated with ^32^P end-labeled linear DNA (198 bp) and 1000-fold excess cold circular DNA and loaded onto a nondenaturing polyacrylamide gel. EV, empty vector; scr siRNA, scrambled siRNA. * band shifted by endogenous Ku heterodimer, ** band shifted by Myc-Ku70 or Myc-Ku80 containing Ku heterodimer. **(B)** Co-immunoprecipitation assays of WCE prepared from cells expressing Myc-tagged Ku80 transgene or EV following Ku80 or scrambled (scr) siRNA knockdown as indicated using Myc pull-down. Antibody against Ku70 was used to detect co-IP of endogenous Ku70 to assess heterodimerization. β-actin was used as loading control for input. Ind, induced; End., endogenous. **(C)** Cells expressing Myc-Ku80 WT, Myc-Ku80 α5 mutant, Myc-Ku80 DEB or EV were subjected to ChIP. Crosslinked cells were lysed, sonicated, and subject to immunoprecipitation with Myc antibody. Telomeric DNA associated with the indicated transgenes was analyzed via dot blot. An Alu probe was used as negative control to monitor association with another repetitive DNA. The lines indicate the removal of irrelevant lanes from the blots. **(D)** and **(E)** Quantification of ChIP experiments as in **C**. The ratio of telomeric DNA **(D)** or Alu **(E)** DNA in IP relative to the respective inputs was evaluated and then for each mutant or EV normalized to Myc-Ku80 WT. Two-way ANOVA and Holm-Sidak’s multiple comparison test were used to determine *p-*value. **, p<0.01; ns, not significant. Error bars indicate standard deviation (SD), n=3. **(F)** ChIP experiments as in **C** but following siRNA mediated depletion of endogenous Ku70 and expression of Myc-Ku70 WT or Myc-Ku70 α5 mutant. EV represents empty vector cell line transfected with scrambled siRNA. **(G)** and **(H)** Quantification of ChIP experiments as in **F** where the ratio of telomeric DNA **(G)** or Alu **(H)** DNA in IP relative to the respective inputs was calculated and normalized to Myc-Ku70 WT. Two-way ANOVA and Holm-Sidak’s multiple comparison test were used to determine *p-*value. **, p<0.01; ns, not significant. Error bars indicate standard deviation (SD), n=4.

To ensure that the lack of DNA end-binding in Myc-Ku80 DEB was not a result of a defect in heterodimerization, we performed co-immunoprecipitation (co-IP) assays and found that the Myc-Ku80 DEB was fully proficient for association with endogenous Ku70 (Fig. 3B). In the co-IP, we observed residual endogenous Ku80 co-immunoprecipitating with Myc-Ku80 WT and Myc-Ku80 α5 mutant but to a lesser degree with Myc-Ku80 DEB (Fig. 3B).

### Ku80 α5, Ku70 α5, and Ku’s DEB function contribute to Ku’s association with telomeres

Having established Myc-Ku80 DEB was impaired for loading onto DNA ends, we used it to further examine whether Ku’s end-binding activity is required for its telomere association. To do this, we performed ChIP assays following induction of Myc-Ku80 WT or Myc-Ku80 DEB mutant expression, but in the absence of knockdown, thus, in the context of endogenous Ku80. This allowed a more stringent evaluation for telomere association, as the Myc-Ku80 DEB would need to compete with endogenous Ku80 for association with the telomere.

As expected, telomeric DNA was enriched in the Myc-Ku80 WT pull down to a much greater extent than Alu DNA. Notably, Myc-Ku80 DEB was significantly impaired for telomere association relative to Myc-Ku80 WT, whereas the relative association with Alu DNA was unchanged (Fig. 3C-E). While Myc-Ku80 DEB was markedly impacted for binding linear DNA in EMSA assays (Fig. 3A), the decrease in telomere association was much less so, and in stark contrast to what had been observed previously with DNA end-binding defective Ku in yeast in which nearly all telomere association was lost (15). This suggests that, as opposed to yeast Ku, where most of Ku’s association with telomeres occurs via end-binding, human Ku associates with telomeres primarily via protein-protein interaction. We also performed ChIP assays to determine if the Ku80 α5 helix also contributes to Ku’s association with telomeres. Notably, despite being proficient for DNA end-binding (Fig. 3A), the Myc-Ku80 α5 mutant also had impaired association with telomeric chromatin (Fig. 3C-E). Thus, perturbation in protein-protein interaction might explain Myc-Ku80 α5 mutant’s reduced telomere association.

We previously implicated Ku70 α5 helix in TRF2 association by demonstrating decreased interaction between the Ku70 α5 mutant and TRF2 in protein-fragment complementation and yeast two-hybrid assays as compared to Ku70 WT (21). To further examine this result, we used ChIP to test whether impaired Ku70 α5 mutant-TRF2 interaction in turn affected Ku70 α5 mutant’s association with telomeres. Like with exogenous Myc-Ku80 WT, telomeric DNA was enriched in the Myc-Ku70 IPs relative to Alu DNA. Interestingly, we found that Myc-Ku70 α5 mutant pulled down significantly less telomeric DNA relative to Myc-Ku70 WT (Fig. 3F-H), implying that the Ku70 α5 helix, which contributes to Ku’s c-NHEJ function and was previously implicated in TRF2 interaction, also supports Ku’s telomere association.

### Myc-Ku80 DEB is impaired for TRF2 interaction whereas Myc-Ku70 α5 mutant shows increased interaction with endogenous TRF2 and its binding partner Rap1

Given the impaired association of the Ku80 α5 mutant and Ku80 DEB with telomeres (Fig. 3F-H), we next assessed if Ku80 α-helix 5 and residues mutated in the Ku80 DEB mutant were involved in interaction with TRF2 via co-IP experiments. We found that Myc-Ku80 α5 mutant’s interaction with immunoprecipitated endogenous TRF2 was comparable to Myc-Ku80 WT (Fig. 4A, B). Surprisingly, however, Myc-Ku80 DEB – TRF2 interaction was significantly decreased relative to Myc-Ku80 WT (Fig. 4A, B). Reciprocal co-IPs, in which Myc-Ku80 was immunoprecipitated, demonstrated similar findings, validating these results (Fig. 4C). One explanation for these results is that the association of Ku and TRF2 in the co-IP experiments was due to a bridging DNA molecule, which would no longer be bound by the Myc-Ku80 DEB. However, the association of Myc-Ku80 nor Myc-Ku70 with TRF2 was unaffected by DNase and Benzonase treatment of the WCEs (Supplementary Fig. S2).

**Figure 4.**
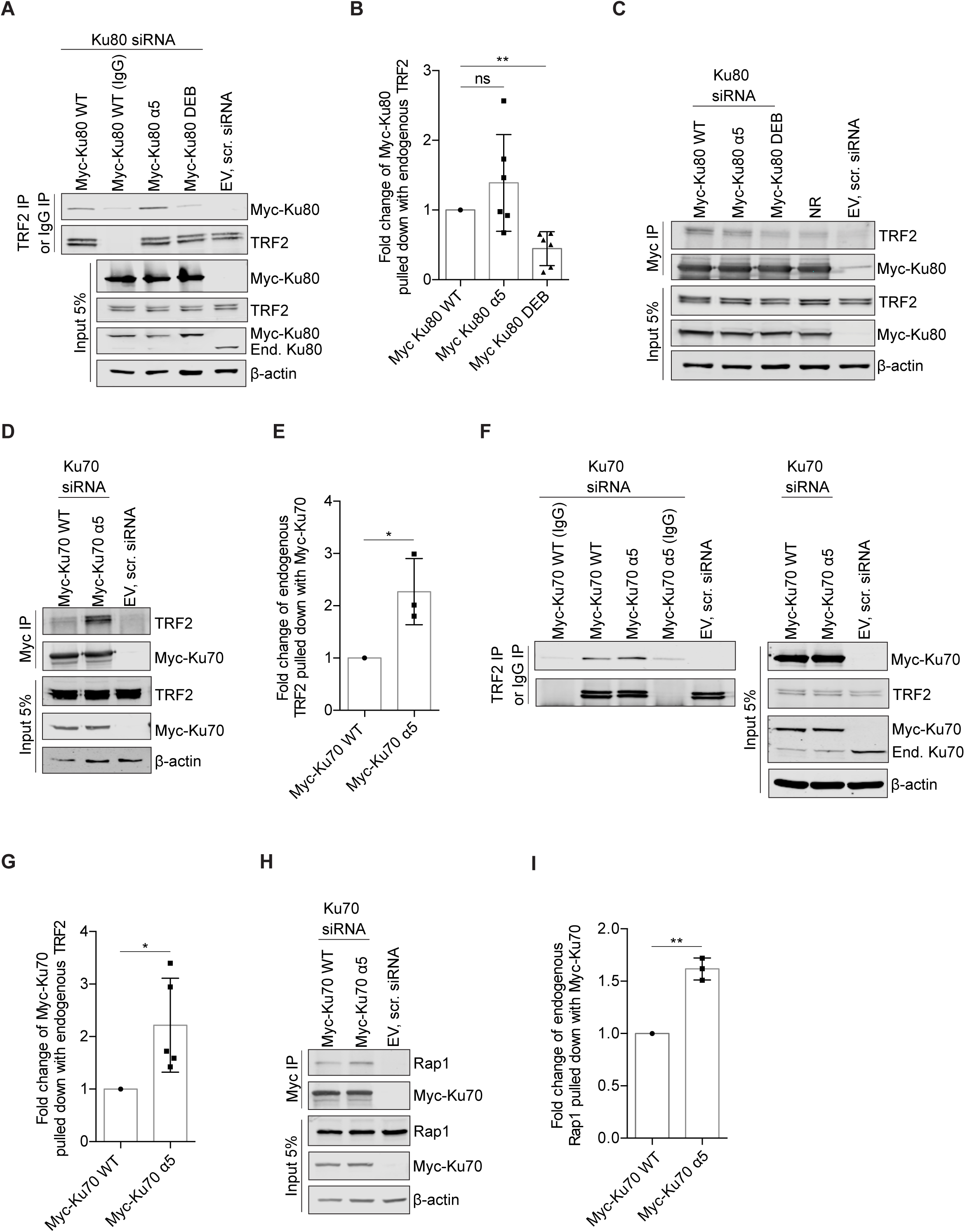
Myc-Ku80 DEB exhibits decreased interaction with TRF2 and Myc-Ku70 α5 mutant exhibits increased interaction with endogenous TRF2 and Rap1. **(A)** Co-IP assays using WCE prepared from cells expressing the indicated Myc-Ku80 transgenes or empty vector (EV) followed by Ku80 or scrambled (scr) siRNA knockdown subject to TRF2 pull-down (IgG control pull-down was performed for Myc-Ku80 WT WCE to monitor background signal). Western blots were analyzed with antibodies against Myc to detect induced (Ind.) Myc-Ku80 and TRF2. β-actin was used as loading control for input. **(B)** Quantification of co-IP experiments as in **A**. The average of the co-immunoprecipitated Myc-Ku80 WT or mutants relative to TRF2 pull-down was determined and normalized to the value for Myc-Ku80 WT. *p-*value was determined by unpaired t-test. **, p<0.01; ns, not significant; error bars indicate SD, n=6. **(C)** Same as **A** except the reciprocal co-IP was performed where WCE were subjected to immunoprecipitation with a Myc antibody. NR indicates nonrelevant lane **(D)** WCE were prepared from cells expressing Myc-Ku70 WT, Myc-Ku70 α5 mutant or EV after siRNA mediated depletion of endogenous Ku70 and subjected to Myc pull-down. Immunoprecipitated proteins were analyzed by western blot with antibodies against Myc and TRF2. **(E)** To quantify co-IPs as shown in **D**, the average of the co-immunoprecipitated TRF2 for Myc-Ku70 WT or Myc-Ku70 α5 mutant relative to Myc pull-down was evaluated and normalized to Myc-Ku70 WT. *p-*value was determined by unpaired t-test. *, p<0.05; error bar indicates SD, n=3. **(F)** Same as **D** except the reciprocal co-IP was performed where WCE were subjected to immunoprecipitation with a TRF2 antibody to pull down endogenous TRF2 or with a control IgG. **(G)** Quantification of co-IPs in **F** and as described in **E.** *p-*value was determined by unpaired t-test. *, p<0.05; error bar indicates SD, n=5. **(H)** WCE similar to **D** were subjected to Myc pull-down and immunoblotted with Myc and Rap1 antibodies. **(I)** Quantification of co-IPs as in **H** where the ratio of co-immunoprecipitated Rap1 relative to Myc pull-down was evaluated and normalized to Myc-Ku70 WT. *p-*value was determined by unpaired t-test. **, p<0.01; error bar indicates standard deviation SD, n=3.

Similar experiments were carried out to investigate the role of Ku70 α5 in TRF2 interaction. In our previous studies, we showed a decrease in interaction between Ku70 α5 mutant and TRF2 proteins in protein fragment complementation assays in which the proteins were fused to split Venus fluorescent protein fragments and transiently overexpressed (21). We probed this finding in our engineered Flp-In T-Rex 293 cell lines via co-IP experiments under conditions where endogenous Ku70 was depleted and Myc-Ku70 WT or Myc-Ku70 α5 mutant was expressed. Unexpectedly, we detected greater association of endogenous TRF2 with immunoprecipitated Myc-Ku70 α5 mutant than with Myc-Ku70 WT (Fig. 4D, E). This increased association was confirmed in reciprocal co-IPs in which endogenous TRF2 was immunoprecipitated (Fig. 4F, G). Since Ku has been shown to interact with Rap1, the shelterin protein that associates with telomeres by binding TRF2, we investigated Myc-Ku70 α5 mutant’s association with Rap1. Similar to the TRF2 result, Myc-Ku70 α5 mutant also exhibited increased association with endogenous Rap1 (Fig. 4H, I). Whether this increase was secondary to increased Myc-Ku70 α5 mutant – TRF2 interaction remains unknown.

### Expression of Myc-Ku80 α5 mutant or Myc-Ku80 DEB affects viability but not telomere length or t-loop integrity in short term cultures

Given Ku’s role in telomere length homeostasis in human cells (26), we sought to determine if cells that express the Ku mutants displayed telomere length defects, which would implicate Ku70 α5, Ku80 α5 or the residues mutated in Ku80 DEB, and, therefore, potentially DNA end-binding, in this essential process. For WT as well as each of the mutants tested, we performed Southern blotting to compare the telomere lengths of cells that had undergone siRNA transfection and doxycycline induction to express transgenes to the respective parental cells that were neither transfected nor induced. Comparison to the respective parental cells was critical as the average telomere lengths of the clonal lines differed. Cells with integrated empty vector (EV) were also used a control. We observed that the mean telomere length of cells expressing Myc-Ku80 WT was slightly less than the uninduced parental line in which the endogenous Ku80 expression was intact (Fig. 5A and Supplemental Fig. S3). Cells that expressed Myc-Ku80 α5 or Myc-Ku80 DEB also showed some decrease in telomere length relative to the corresponding uninduced cells, although this effect was variable for Myc-Ku80 DEB. In cells induced to express Myc-Ku70 WT or Myc-Ku70 α5 mutant, telomere lengths remained more or less comparable to the respective parental lines (Fig. 5B). Slight telomere shortening was observed for cells with an integrated empty vector that were treated with doxycycline and transfected with scrambled siRNA. Taken together, we did not see differences in telomere lengths that could be attributed to the Ku mutants. Moreover, unexpectedly, the mean telomere length of the EV cell line depleted of endogenous Ku80 was comparable to the EV parental line. Similarly, knockdown of endogenous Ku70 did not impact the telomere length of the EV cell line (Fig. 5B). One explanation for the absence of telomere shortening in these Ku80- and Ku70-depleted lines, as noticed previously (26,34), is that there was insufficient time to observe a difference in telomere length in this system.

**Figure 5.**
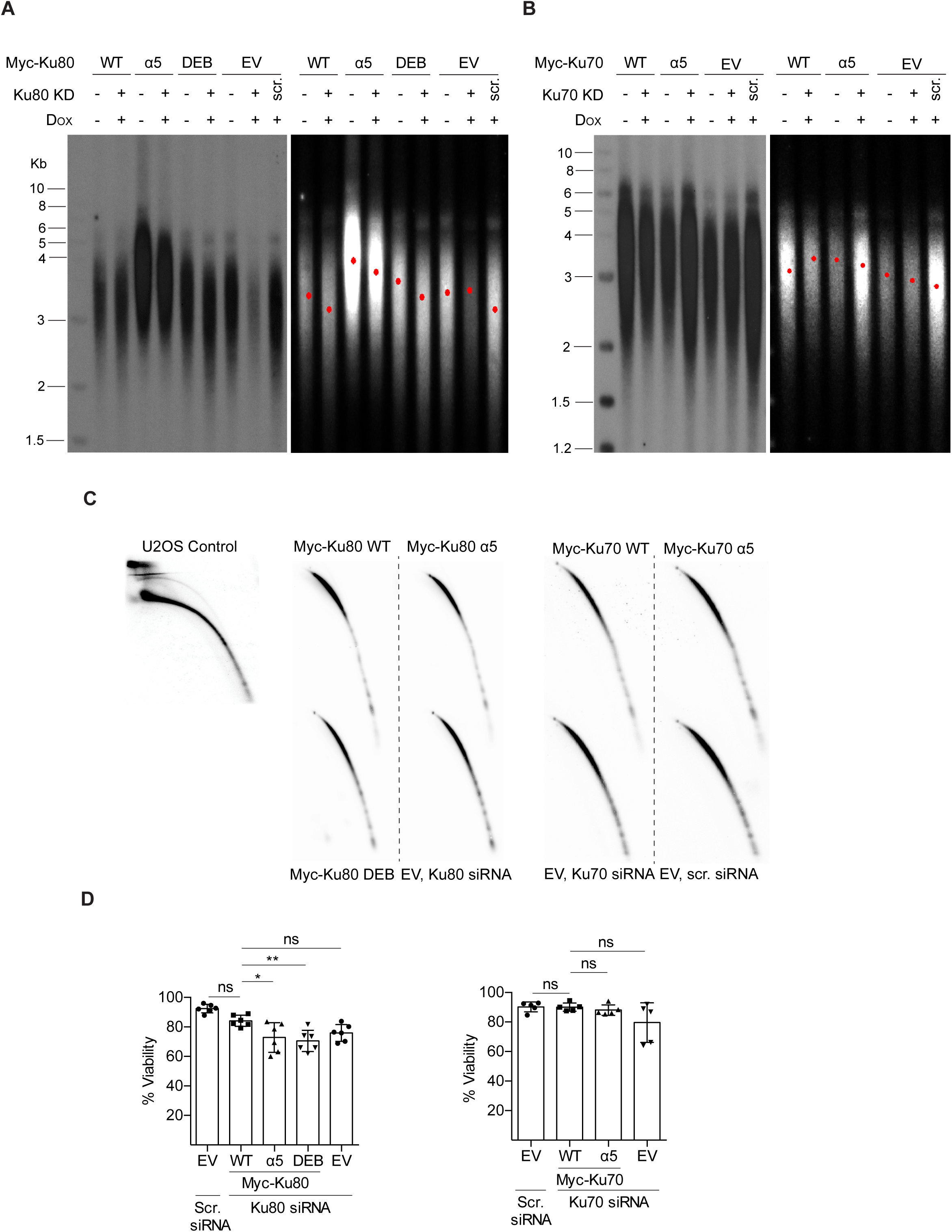
Myc-Ku80 α5 mutant or Myc-Ku80 DEB expression compromises cell viability but does not lead to telomere shortening or t-circle formation in short term cultures. **(A)** Representative Southern blot analysis to measure telomere lengths of indicated cell lines uninduced or induced to express Myc-Ku80 transgenes. Empty vector cell line transfected with scrambled (scr) or Ku80 siRNA was used as positive and negative controls. KD indicates siRNA-mediated knockdown. Telomere Southern blots were quantified via TeloTool software and the red dot in each lane denotes mean telomere length. Additional representative experiment included in Supplemental Figure S3 **(B)** Same as **A** except telomere lengths of indicated cell lines uninduced or induced to express Myc-Ku70 transgenes. Additional representative experiment included in Supplemental Figure S3 **(C)** 2D gel electrophoresis of HinfI/RsaI digested genomic DNA prepared from indicated cell lines that were subjected to either Ku70 or Ku80 knockdown. U2OS control cell line showing t-circles **(D)** Analysis of cell viability by Annexin V staining of indicated cell lines expressing Myc-Ku80 or Myc-Ku70 transgenes. Empty vector cell line transfected with scrambled (scr) or Ku70 or Ku80 siRNA was used as controls.

A consequence of loss of Ku80 in the human HCT116 colorectal carcinoma cell line is the appearance of extrachromosomal telomeric circles, known as t-circles, which are thought to be produced by the improper processing of telomeric t-loops by HR (26). To monitor t-circles, we used 2D gel electrophoresis and found that expression of Myc-Ku80 α5, Myc-Ku80 DEB or Myc-Ku70 α5 mutant did not result in t-circle formation (Fig. 5C). Moreover, consistent with the Southern blot results, we did not detect t-circles in the EV cell line that was subjected to either Ku70 or Ku80 knockdown, indicating the absence of telomere loss upon depletion of Ku in this experimental system (Fig. 5C).

Studies have demonstrated Ku to be an essential gene in human cells (32,39). To address if expression of Ku mutants compromised cell viability, we performed annexin V assays. While the percentage of viable cells in Myc-Ku80 WT did not vary significantly from the control EV line transfected with scrambled siRNA, we noticed a significant reduction in cell viability in cells expressing Myc-Ku80 α5 mutant and Myc-Ku80 DEB (Fig. 5D). Expression of the Myc-Ku70 α5 mutant, however, did not affect cell viability.

## DISCUSSION

An intriguing conundrum in telomere biology is how the DNA end-binding Ku heterodimer associates with functional telomeres to exert its telomeric functions without also initiating c-NHEJ. We aimed to address this question using cen3tel cell lines with different telomere lengths and potential separation-of-function human Ku mutants that impact either Ku’s end-binding function or its ability to interact with its protein binding partners. Additionally, we sought to understand if the functional and structural organization of Ku identified in yeast was conserved in human cells, where Ku has been shown to be essential for its function in telomere maintenance (26,34).

The results of the ChIP experiments to interrogate Ku’s localization to telomeres in cen3tel cells are most consistent with the model of Ku associating with human telomeres largely via protein-protein interaction (Fig. 1). Further evidence for this finding emerged from probing the DNA end-binding defective Ku80 DEB’s telomere association in Flp-In T-REx 293 cells, which showed some reduction in its ability to localize to telomeres, but a considerable amount of telomeric DNA still immunoprecipitated with Ku80 DEB despite the mutant protein displaying a marked impact on DNA end association (Fig. 3A and 3C-E). Moreover, Ku80-DEB was also impaired for TRF2 interaction (Fig. 4A-C), which may have contributed to the reduction in telomere association rather than a defect in DNA end-binding. Overall, these data demonstrate a striking difference in Ku’s mechanism of association with telomeres between yeast and humans. In contrast to yeast where DNA binding was demonstrated to be the primary mode by which Ku associates with telomeres (15), human Ku’s recruitment to telomeres is mainly mediated through protein-protein interactions, presumably via its associations with shelterin components such as TRF2 and possibly yet to be identified telomere binding partners. How these specific protein interactions influence Ku’s functions at human telomeres remains to be understood.

Meanwhile, it remains an open question if some of Ku’s association with telomeres in human cells also occurs through direct engagement with the chromosomal ends. It is possible that Ku’s localization to human telomeres via end-binding is confined to a specific phase of the cell cycle, such as S phase, when the t-loop presumably disassembles to facilitate telomere replication. Alternatively, far fewer Ku molecules might engage telomeres via end-binding compared to recruitment through protein-protein interactions and, thus, influence the outcome of the cen3tel ChIP experiments to a lesser extent.

Our goal was to generate a separation-of-function human Ku80 DEB akin to the Ku DNA end-binding mutant we had previously characterized in yeast, which was defective for end-binding but proficient for interactions with its telomere binding partners, specifically Sir4 and TLC1 (15). Surprisingly, the human Ku80 DEB was defective for both telomere association and TRF2 interaction (Fig. 3C-E and Fig. 4A-C), suggesting either shared residues on Ku80 for DNA end-binding and TRF2 interaction or that Ku has to load onto telomeric DNA to interact with TRF2. However, neither Ku70-TRF2 nor Ku80-TRF2 association in co-IP experiments were impacted by DNase/Benzonase treatment, suggesting DNA binding is not required. If TRF2 interacts with Ku via residues also used for DNA end-binding, this might be a mechanism to prevent Ku from loading onto the telomeric DNA end and, thereby, suppress Ku’s c-NHEJ functions at telomeres. Alternatively, it is possible that the mutations introduced to generate Ku80 DEB led to a conformational change of the protein, affecting Ku’s interaction with TRF2.

We investigated the Ku80 α5 helix in human cells, since the comparable helix was previously implicated for its role in telomeric silencing in budding yeast (19). Our results indicate Ku80 α5 mutant is impaired for telomere association (Fig. 3C-E), which is notable given humans lack an ortholog of the Sir4 protein which binds the Ku80 α5 helix in yeast. However, the defect in telomere association is neither due to deficiencies in DNA end-binding or TRF2 interaction (Fig. 4A-C), but might be attributed to impaired interaction with TRF1 or Rap1 or some other telomere associated factor. These data further underscore Ku’s association along the telomeric DNA tract by protein-protein interactions.

Previous studies had established a role for Ku70 α5 helix in c-NHEJ in yeast and mammalian cells (19-21). We set forth to test Ku70 α5 mutant’s telomere association and found that, although it retained Ku’s DNA end-binding property (Fig. 3A), the Ku70 α5 mutant was impacted for telomere association (Fig. 3F-H). However, unexpectedly, we detected increased interaction between Ku70 α5 mutant and endogenous TRF2 and Rap1 (Fig. 4D-I). Why then do we observe less Ku70 α5 mutant at telomeres? We speculate that Ku70 α5 mutant’s increased association with TRF2 might be occurring at DSBs, which could sterically hinder Ku’s association with a DNA repair factor and, thus, contribute to the c-NHEJ defect.

Cells expressing Ku80 α5 mutant and Ku80 DEB were impaired for viability while expression of Ku70 α5 mutant did not affect cell survival (Fig. 5D). Moreover, depletion of Ku70 or Ku80 in the EV cell line did not impact the viability of the cells, implying that the remaining endogenous protein levels following knockdown was sufficient to sustain the survival of the cells. Expression of the Ku80 α5 mutant or Ku80 DEB, however, might harm the functionality of the remnant endogenous protein, thus, affecting cell viability. It remains to be explored if the decreased viability is a consequence of telomere dysfunction in human cells. While we did not detect any changes in telomere length or release of t-circles at 4 days following depletion of endogenous Ku, longer term studies might be required to examine if expression of the Ku mutants lead to telomere length defects.

## SUPPLEMENTARY DATA

Supplementary Data are available at NAR online.

