## Supplementary material for "The DNA end-binding protein Ku associates with human telomeres primarily via protein-protein interactions": Supplentary Figures S1-S3

**Supplementary Figure S1 Ku associates with telomeres via protein-protein interactions.** **(A)** Additional representative ChIP of cen3tel cells corresponding to a different set of PDs and approximate telomere lengths as indicated. After crosslinking with formaldehyde, cells were sonicated, and subjected to lysis followed by immunoprecipitation of endogenous Ku80 with a Ku80 antibody. Telomeric and Alu DNA associated with the respective PDs was analyzed via slot blot. IgG was used as negative control and Alu DNA was used as repetitive DNA control. **(B)** Table showing the percentage of telomeric DNA immunoprecipitated with Ku80 or control IgG relative to telomere input and Alu input for the respective PDs.

**Supplementary Figure S2 The association of Myc-Ku80 or Myc-Ku70 with TRF2 is DNA independent. (A)** Co-IP experiments with WCE isolated from cells expressing Myc-Ku80 WT after siRNA knockdown of endogenous Ku80. The lysates were subjected to DNase and Benzonase treatment, where indicated, followed by immunoprecipitation with a TRF2 antibody (IgG pull-down was used as control to account for background signal). For the western blot, antibodies against Myc and TRF2 were used to detect Myc-Ku80 and TRF2, respectively. β-actin was used as loading control for input. **(B)** Same as **(A)** except WCE were prepared from cells expressing Myc-Ku70 WT following siRNA mediated depletion of endogenous Ku70. Antibody against Myc was used to observe Myc-Ku70 signal in the western blot.

**Supplementary Figure S3 Expression of Ku mutants or depletion of endogenous Ku does not lead to telomere shortening. (A)** Southern blot analysis to monitor differences in telomere lengths of indicated cell lines uninduced or induced to express Myc-Ku80 transgenes. Empty vector cell line transfected with scrambled (scr) or Ku80 siRNA was used as positive and negative controls. TeloTool software was used to quantify telomere Southern blots and the red dot in each lane represents mean telomere length. **(B)** Same as **A** except telomere lengths of indicated cell lines uninduced or induced to express Myc-Ku70 transgenes.

Supplementary figure 1

**A**

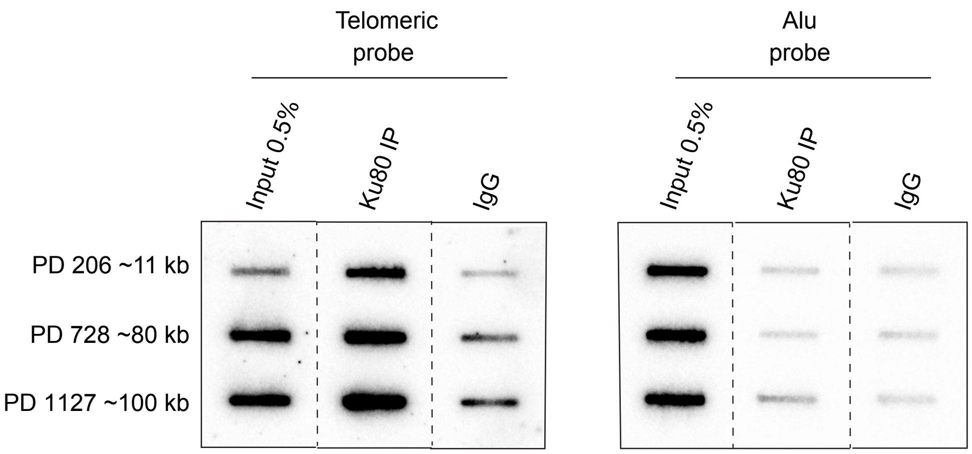


**B**

| | Telo IP/ Alu input | | Telo IP/ Telo input | |
|---|---|---|---|---|
| | Ku80 IP | IgG IP | Ku80 IP | IgG IP |
| PD 206 | 0.37 | 0.04 | 1.53 | 0.18 |
| PD 728 | 0.79 | 0.11 | 1.03 | 0.14 |
| PD 1127 | 1.28 | 0.15 | 1.30 | 0.16 |

Supplementary Figure S2

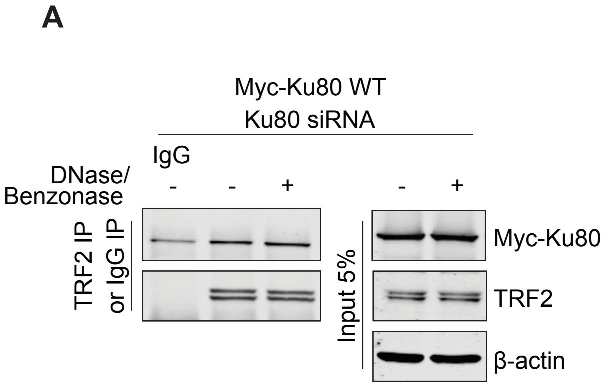

Supplementary Figure S3

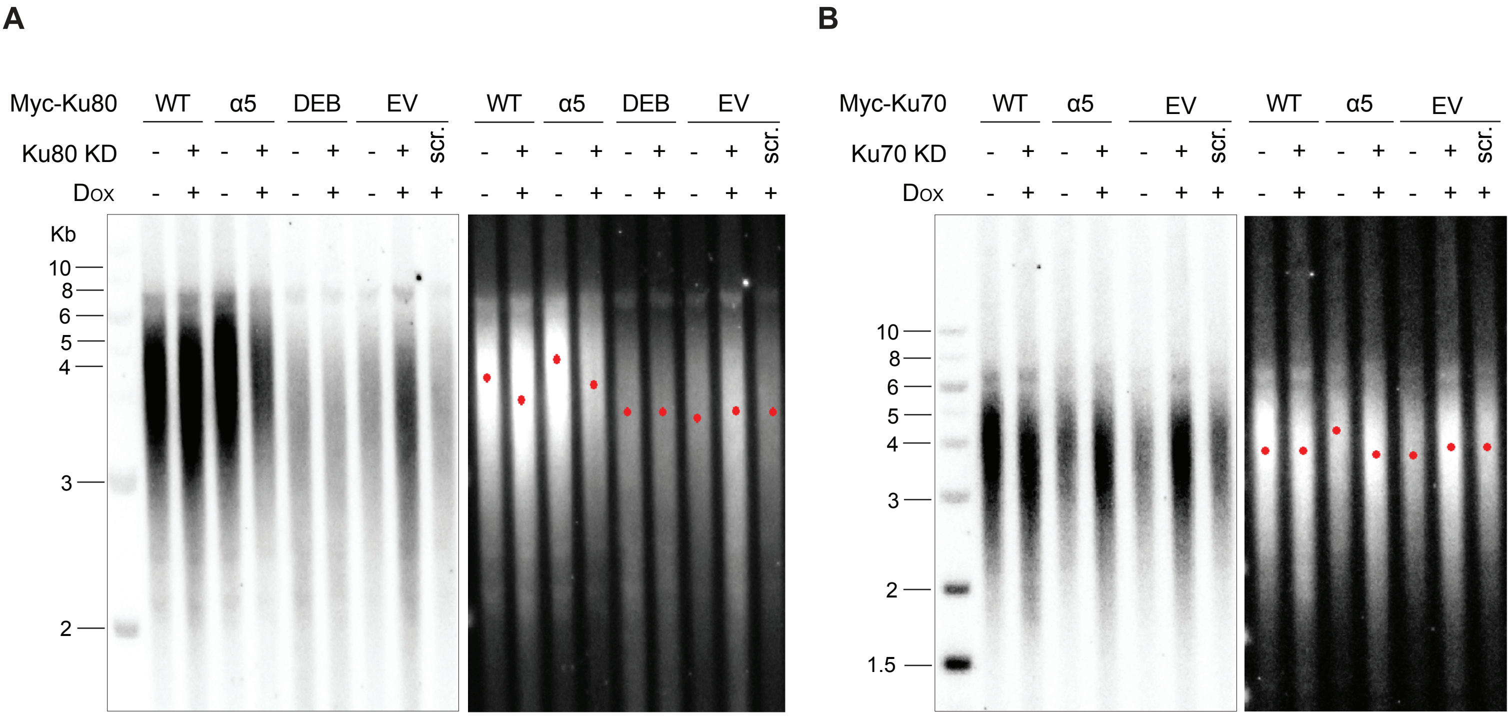
